# Enigmatic Diphyllatea eukaryotes: Culturing and targeted PacBio RS amplicon sequencing reveals a higher order taxonomic diversity and global distribution

**DOI:** 10.1101/199125

**Authors:** J.S. Orr Russell, Zhao Sen, Klaveness Dag, Yabuki Akinori, Ikeda Keiji, Makoto M. Watanabe, Shalchian-Tabrizi Kamran

## Abstract

Diphyllatea is an ancient and enigmatic lineage of unicellular eukaryotes that possesses morphological features common to other deeply diverging eukaryotes, such as Amoebozoa and Excavata. In reconstruction of the evolutionary processes underlying diversification and morphological innovation among eukaryotes, Diphyllatea plays a key role together with other orphan lineages. Despite being of evolutionary significance, only three species of Diphyllatea have descripted morphology, with molecular data available from fewer. The lack of data means that the actual diversity of this key lineage of eukaryotes remains unresolved. We here present a first attempt to understand the species diversity and higher order structure of the Diphyllatea phylogeny. We have cultured several new strains, described these morphologically, and amplified their rRNA. We have sampled DNA from multiple globally distributed sites, using these as templates in a Diphyllatea-specific PCR. In contrast to recent diversity studies, which use short variable gene regions, we amplify nearly the whole 18S rRNA gene, and sequence using PacBio RS II technology, to provide enough information to resolve historically ancient speciation events. Phylogenetic inference of Diphyllatea rRNA reveals three deeply branching and distinct clades of Diphyllatea, here named Diphy I – III. Diphy I and II include the genera *Diphylleia* and *Collodictyon*. Notably, Diphy III is here shown as novel phylogenetic clade with all strains investigated having a congruent morphology to *Collodictyon triciliatum* (Diphy II). Altogether, Diphyllatea seems to constitute two morphotypes, a biflagellate (i.e. Diphy I) and a quadraflagellate (i.e. Diphy II and III) form, congruent with earlier descriptions of *Diphylleia and Collodictyon.* Further, our targeted environmental sequencing approach, which includes specific PCR primers, reveals a wider global distribution of Diphyllatea than earlier known. Altogether, the described protocol shows the usefulness of combining long amplicon high-throughput sequencing and lineage-specific PCR approach in surveys of enigmatic eukaryote lineages.

## Introduction

The class Diphyllatea is a group of protists that holds a deep and distinct position in the eukaryote tree, most likely as one of the earliest diverging lineages (Zhao et al., 2012; Brown et al., 2013; Zhao et al., 2013; Burki et al., 2016). Presently, only a few species are described using traditional microscopic methods. Initially, the class Diphyllatea and the order Diphylleida were proposed to encompass the biflagellate *Diphylleia rotans* and the quadraflagellate *Collodictyon triciliatum* (Brugerolle et al., 2002). However, a recent revision of the systematic classification changed this class to include the species *Sulcomonas lacustris* and several older synonyms (e.g. *C. sparseovacuolatum* for *C. triciliatum* and *Aulacomonas submarina* for *D. rotans*) (Brugerolle and Patterson, 1990; Brugerolle, 2006). Currently, based on morphological features, with only 18S rRNA provided from *D. rotans*, Diphyllatea is proposed to consist of the three genera *Collodictyon*, *Diphylleia* and *Sulcomonas* (Cavalier-Smith, 2013; Ruggiero et al., 2015), with the two prior constituting the family Diphylleida and the former family Sulcomonadidae. The three representative species (*C. triciliatum*, *D. rotans* and *S. lacustris*) have been previously investigated by light and electron microscopy (Carter, 1865; Francé, 1899; Massart, 1920; Klaveness, 1995; Bo-Ra et al., 2006; Brugerolle, 2006; Mohamed and Al-Shehri, 2013). They share a heart or egg shape and possess a ventral groove (Francé, 1899; Wawrik, 1973; Klaveness, 1995), more or less dividing the body longitudinally, but the size range of the identified species is variable (15-60 μm length for *Collodictyon,* 20-25 μm length for *Diphylleia* and 8-20 μm length for *Sulcomonas*).

As the description of species diversity and the erection of the whole taxonomic unit of Diphyllatea were based on microscopic observations, one could expect that sequencing surveys of environmental DNA would detect a larger number of cryptic species. On the contrary, excluding the original *D. rotans* 18S rRNA (AF420478) from Brugerolle et al. 2002, only a single partial 18S rRNA sequence of Diphyllatea-like organisms (DLOs) has been reported from a Tibetan freshwater lake (AM709512), and until now, no DLOs have been classified from other water samples (Wu et al., 2009). It is known that environmental PCR with the use of group-specific primers can effectively amplify the diversity of some unicellular eukaryotes (Bass and Cavalier-Smith, 2004; Potvin and Lovejoy, 2009; Bråte et al., 2010), but such an approach has never before been applied to Diphyllatea. Hence, its diversity may currently be underestimated. Accordingly, the geographical distribution of the class, based on molecular data, is presently limited to China, France and Norway, with lower resolution at the family and genus level (Brugerolle, 2006; Wu et al., 2009; Zhao et al., 2012). Though, morphological data seemingly suggests a possible global distribution of *C. triciliatum* (Carter, 1865; Rhodes, 1917; Lackey, 1942; Klaveness, 1995; Sánchez et al., 1998).

Thus, the objective of this study is to investigate possible cryptic diversity and distribution of Diphyllatea by firstly studying the morphology of novel cultured strains, and secondly by the amplification of DLO rRNA from marine and freshwater samples. Additional database mining will allow for the confirmation of the classes diversity and distribution.

Recently, eukaryotic diversity studies have used Illumina (predominantly the MiSeq platform) for sequencing rRNA amplicons. As Illumina has a restricted read length, diversity studies have been limited to an amplicon maximum of approximately 450bp. A result of this being that studies have either focused on short hypervariable regions of 18S rRNA (Hugerth et al., 2014; de Vargas et al., 2015; Bradley et al., 2016), ITS (Taylor et al., 2016; Al-Bulushi et al., 2017), or 28s rRNA (Asemaninejad et al., 2016; Mueller et al., 2016). These regions, despite being variable, sometimes lack enough sequence variation to be able to divide some genera to the species level. Further, focusing on separate, non-overlapping rRNA regions makes it difficult to study amplicons in a comparative phylogenetic context. Conversely, the study of long amplicons has traditionally involved cloning and sanger sequencing (Edgcomb et al., 2011; Thomas et al., 2012; Bachy et al., 2013), a time consuming and costly method when high depth is desired. The Pacific Bioscience (PacBio) RS sequencing platform offers an alternative to short Illumina reads by providing long (>20kb) sequencing reads. As such PacBio RS may also represent an alternative to cloning and sanger sequencing for longer rRNA amplicons. However, unlike Illumina, that provides high-output and high-quality reads, PacBio SMRT (Single Molecule Real Time) cells have a lower output and higher error rate (∼15% with the P4-C2 chemistry) due to the random addition of incorrect nucleotides (Eid et al, 2009; Koren et al, 2012). To overcome the random error rate, DNA is ligated into SMRT bells; circular DNA fragments that allow multiple sequence passes, a process termed Circular Consensus Sequencing (CCS) (Travers et al., 2010). To date, PacBio RS has mainly been applied to genome and more recently transcriptome sequencing (Hoang et al., 2017; Kuo et al., 2017; Orr et al., 2017b, a). Though, a few studies have shown the platforms viability for studying 16s rRNA diversity of prokaryotes (Fichot and Norman, 2013; Mosher et al., 2013), and more recently eukaryotic rRNA amplicons (Jones and Kustka, 2017; Tedersoo et al., 2017). Jones and Kustka sequenced the V7-9 region of 18S rRNA (454-615bp) to answer questions about total eukaryotic diversity from marine samples (Jones and Kustka, 2017). Tedersoo et al., targeted the V4 (18S)-D3 (28S) region focusing on total eukaryotic and fungal diversity from soil samples, confirming PacBio as an alternative for metabarcoding of organisms with low diversity for reliable identification and phylogenetic approaches (Tedersoo et al., 2017). To date, no studies have applied PacBio to sequence targeted 18S rRNA amplicons of lengths >1000bp.

For this reasoning, the secondary objective of this study is to assess the PacBio RS platform as an alternative to traditional cloning and Sanger sequencing methods for the study of long environmental amplicons; in particular, >1000bp targeted 18S rRNA amplicons from environmental DLOs.

## Materials and Methods

### Culture isolation and maintenance

Asian strains of DLO were established by a single cell isolation method from localities in Japan, Thailand and Vietnam (Table 1). The isolated strains from Asia were inoculated into the freshwater medium URO (Kimura and Ishida, 1985) with endogenous cyanobacteria (*Microcystis*, strain no. NIES-44) as food and established as cultures. The investigated Norwegian strain of *Collodictyon triciliatum* (i.e. strain Å85) was a clonal isolate (from a single cell) from Lake Årungen and was initially cultured on WC-medium (Guillard and Lorenzen, 1972) with the cryptomonad *Plagioselmis nannoplanktica* or a strain of the green algae *Chlorella* as food (Klaveness, 1995). Subsequently, all cultures were kept in BG11 1/2 medium (Stanier et al., 1971), with *Microcystis* strain CYA-43 provided by the Norwegian Institute of Water Research (NIVA–www.niva.no). All cultures were grown using the following conditions: 17 °C, 250 μMol m^-1^ sec^-1^ of daylight-type fluorescent light at a 14 / 10 (L / D) cycle.

**Table 1.** Sampling locations for cultured Diphyllatea-like organisms and environmental DNA: Environmental DNA samples were chosen based on size fractions encompassing the known cell size of Diphyllatea species / strains: 8-60μm (see Fig. 1 and Brugerolle et al 2002, Brugerolle 2006). All environmental DNA was sampled subsurface.

| Strain/sample | Sampling locality |
| --- | --- |
| **Cultured samples** |  |
| Å85 | Lake Årungen, Ås, Norway (59°41'N 10°44'E) |
| KIINB | Lake Inba, Thiba, Japan (35°44'N 140°10'E) |
| KIKNR01, 02, 03 | Kaen Nakon Reservoir, Thailand (16°24'N 102°50'E) |
| KIVT01, 02, 03, 04 | Hồ Dầu Tiếng, Vietnam (11°23'N 106°17'E) |
| KIVTT01, 02 | Turtle Farm, Ha Tinh, Vietnam (18°19'N 105°53'E) |
| **Freshwater DNA** |  |
| Årungen | Lake Årungen, Ås, Norway (59°41'N 10°44'E) |
| BOR41 | Lake by Kinabatangan river, Borneo, Malaysia (5°25'N 117°56'E) |
| BOR42 | Pond A, Sandakan, Borneo, Malaysia (5°50'N 118°7'E) |
| BOR43 | Pond B, Sandakan, Borneo, Malaysia (5°50'N 118°7'E) |
| LD_DERW20 | Derwent water, UK (54°34'N 3°8'W) |
| LD_ESTH20 | Esthwaite water, UK (54°21'N 2°59'W) |
| LD_BASS2, 20 | Bassenthwaite lake, UK (54°40'N 3°13'W) |
| SA78, 81 | Pond, Cape Town, South Africa (33°56'S 18°24'E) |
| **Marine DNA** |  |
| NB038 | Naples Bay, Italy (40°49'N 14°18'E) |
| RA119 | Roscoff, France (48°43'N 4°2'W) |
| VA105 | Varna, Bulgaria (43°11'N 28°0'E) |
| 20F268 | Oslo Fjord, Norway (59°27'N 10°32'E) |

### Microscopy

Light microscopy of the 11 DLOs was conducted using a Nikon Diaphot inverted microscope. Differential interference contrast (DIC) micrographs and video of DLO cells was conducted using a Nikon D-series digital camera (D1 and D300S) connected to a TV screen for focusing on, and following motile cells. Electron microscopy (EM) was done by the negative staining of whole cells, after drop fixation on grids by osmium vapour (Klaveness, 1995).

### DNA isolation, PCR and sequencing

DNA was isolated from 50ml of each culture by pelleting cells by centrifugation at 1500 rpm and 4°C for five minutes, followed by standard CTAB chloroform / isoamylalcohol extraction and subsequent ethanol precipitation (Doyle and Doyle, 1987). A ∼6.3kb region of the rRNA operon, covering the 18S, ITS1, 5.8s, ITS2 and 28s regions, was amplified as one continuous fragment with the forward primer NSF83 and the reverse primer LR11 (Table 2) utilizing Phusion High-Fidelity DNA polymerase, 35 cycles and a 50°C annealing (ThermoFisher). The single ∼6.3kb PCR products were cleaned using Chargeswitch PCR Clean-up kit (ThermoFisher) and then Sanger sequenced (GATC Biotech, Germany) as separate fragments, utilizing primers outlined in Supplementary Table 2. Additional sequencing primers were designed using Primaclade (Gadberry et al., 2005). The separate fragments were subsequently quality checked and assembled using the Phred / Phrap / Consed package (Gordon et al., 1998) under default settings. Additional manual editing of the 11 contigs was performed in Mesquite v3.1 (Maddison and Maddison, 2017).

**Table 2.**
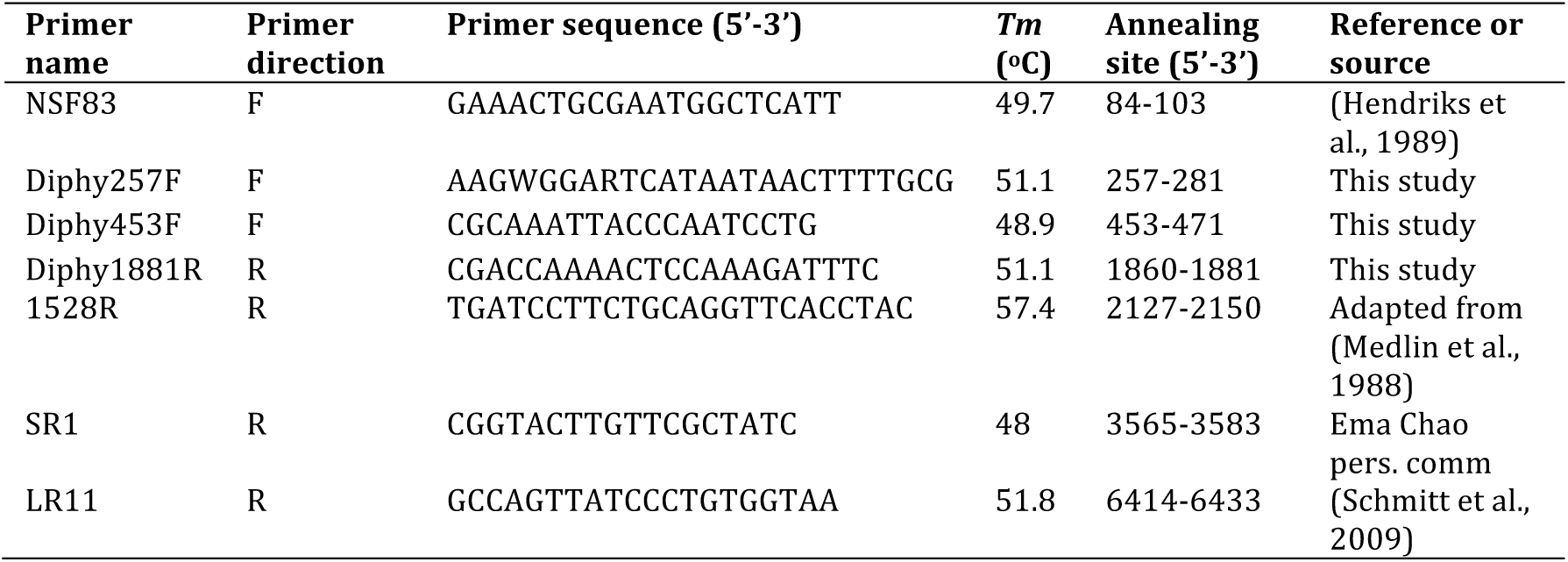
List of primers used in this study: Primer annealing site is based on *Collodictyon* KIVT02 sequence, start is 83bp prior to account for NSF83s annealing site. *Tm* is calculated using oligocalc (Kibbe, 2007). Primers used for Sanger sequencing are listed in Supplementary Table 1.

### Primer design and specificity confirmation

For optimal primer design with high specificity to the Diphyllatea clade, all available orthologous DLO sequences were used for alignment construction; Collo Å85 and KIVT02 rRNA were used as blastn queries to extract DLO sequences deposited in the NCBInr database (http://blast.ncbi.nlm.gov/Blast.cgi) using default parameters. Further, *Diphylleia rotans* NIES-3764 rRNA (isolated in Amakubo, Ibaraki prefecture, Japan), taken from an unconnected genome project, was additionally used as a query and included in subsequent analyses. The resulting sequences were aligned together with the 11 culture sequences using the MAFFT Q-INS-I model (Katoh and Standley, 2013), considering secondary RNA structure (default parameters used). The alignment was then manually checked and edited using Mesquite v3.1 (Maddison and Maddison, 2017) before designing primers with Primaclade (Gadberry et al., 2005). All potential primers were tested for specificity to the Diphyllatea clade by checking sequence identity against non-Diphyllatea sequences in the silva rRNA database in addition to an rRNA alignment with a broad sample of eukaryotic taxa (Cavalier-Smith and Chao, 2010). OligoCalc (Kibbe, 2007) was applied to check for self-complementarity and calculate primer *Tm*. The Diphyllatea specific primers with highest potential were then utilized in PCRs to confirm amplification of Diphyllatea rRNA, with optimal annealing temperature being established, and the non-amplification of DNA template external to Diphyllatea (a broad DNA mix from multiple cultures held in our lab). The Diphyllatea specific rRNA primers designed in this study are listed in Table 2.

### Environmental DNA and confirmation of Diphyllatea

Environmental DNA was sampled from Lake Årungen by collecting and filtering two liters of surface water through a Whatman GF / C glass-fiber filter with pore sizes of approximately 1 μm. DNA was isolated as previously outlined. Dr. David Bass (NHM) kindly provided supplementary freshwater DNA samples (Table 1) from Borneo, South Africa, and the UK. Dr. Bente Edvardsen (UiO) in collaboration with BioMarKs (Logares et al.) kindly provided marine DNA samples (Table 1) from Bulgaria, France, Italy, and Norway. Eukaryotic DNA was confirmed for all samples by PCR, as previous, with a 55°C annealing, using the universal 18S rRNA primers NSF83 and 1528R (Table 2). Diphyllatea clade specific PCR was subsequently performed on all environmental DNA samples targeting the 18S rRNA region with the primers Diphy257F and Diphy1881R (∼ 1624bp: see Table 2), as previous, with a 55°C annealing temperature. Additionally, the annealing temperature for the Diphyllatea clade specific PCR (Diphy257F-Diphy1881R) was lowered by 5°C to allow primers to anneal to possible novel DLO template rRNA with lower sequence identity. Finally, for those environmental templates that gave no PCR-product with the above primer-pair, a pair with a lower specificity to the Diphyllatea clade was employed; Diphy453F and 1528R (∼1697bp: see Table 2) amplifies a range of eukaryotes including Diphyllatea. PCR was as previous with a 55°C annealing temperature. A positive (Å85 and KIVT02 DNA) and negative control were employed for all PCRs. Positive amplicons were cleaned using Chargeswitch PCR Clean-up kit (ThermoFisher) or the Wizard SV gel and PCR clean-up system (Promega) and used for down-stream processing.

### PacBio barcodes, library prep and amplicon sequencing

As a more economical and efficient alternative to cloning, PacBio RS II was employed to achieve higher sequencing depth of the long environmental rRNA amplicons. PCR primers with symmetric (reverse complement) PacBio barcodes (21bp) were attached to the separate rRNA amplicons by PCR: a 2μl 1:10 dilution of template DNA (rRNA amplicon) was used as input in a two-step PCR protocol using Phusion High-Fidelity DNA polymerase (ThermoFisher), with a 72°C 90 second annealing and 20 cycles. The resulting PCR product was cleaned, as previous, before successful attachment of PacBio barcodes was confirmed using Bioanalyzer (Agilent Technologies); comparing product length pre-and post-barcode PCR. A single SMRTcell was prepared and sequenced, multiplexing both the Diphy257F-Diphy1881R (∼1624bp) and the Diphy453F-1528R (∼1697bp) amplicons (Table 3 & Supplementary Table 2). Input concentrations for each sample were adjusted, dependent on quantity. The Norwegian Sequencing Centre (NSC), Oslo, Norway, performed library preparation and sequencing. The Library was prepared using Pacific Biosciences 2 kb library preparation protocol, before sequencing with the PacBio RS II instrument using P4-C2 chemistry. Filtering was performed using Reads of Insert protocol on SMRT portal (SMRTAnalysis v2.2.0.p1 build 134282). Default settings (Minimum number of passes=1 and Minimum Predicted Accuracy=0.9) were used.

**Table 3.**
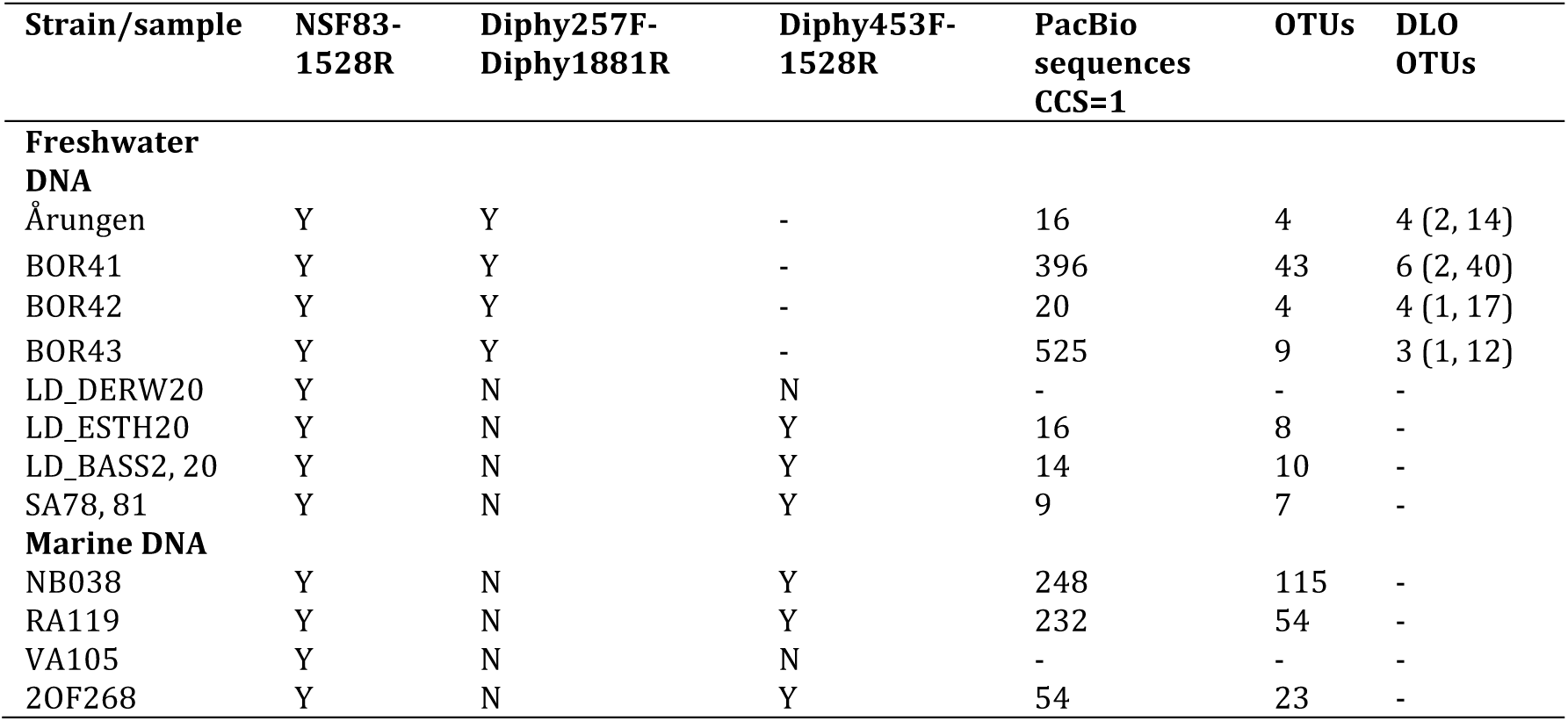
PCR and sequencing results for environmental amplicons: Y = PCR product, N = no PCR product. The value for the PacBio reads (column five) is minus chimeras identified in Uchime. For DLO OTUs (last column) the number in brackets represents firstly the number of OTUs constituting >1 read, and secondly the total number of reads these non-unique OTUs constitute; these are represented in the rRNA phylogeny (Figure 3 and Supplementary Figure 2). For additional sequencing results per PacBio barcode see Supplementary Table 2. A complete BLAST result against NCBInr for all 281 OTUs is provided as supplementary data. Though briefly, and outside the scope of this study, for the marine samples (192 OTUs): bivalve, ciliate, diatom, dinoflagellate and segmented worm rRNA were most abundant. For the freshwater samples (25 OTUs): ciliate, cryptomonad and diatom rRNA were most common

| Strain/sample | NSF83-1528R | Diphy257F-Diphy1881R | Diphy453F-1528R | PacBio sequences CCS=1 | OTUs | DLO OTUs |
| --- | --- | --- | --- | --- | --- | --- |
| **Freshwater DNA** |  |  |  |  |  |  |
| Årungen | Y | Y | - | 16 | 4 | 4 (2, 14) |
| BOR41 | Y | Y | - | 396 | 43 | 6 (2, 40) |
| BOR42 | Y | Y | - | 20 | 4 | 4 (1, 17) |
| BOR43 | Y | Y | - | 525 | 9 | 3 (1, 12) |
| LD_DERW20 | Y | N | N | - | - | - |
| LD_ESTH20 | Y | N | Y | 16 | 8 | - |
| LD_BASS2, 20 | Y | N | Y | 14 | 10 | - |
| SA78, 81 | Y | N | Y | 9 | 7 | - |
| **Marine DNA** |  |  |  |  |  |  |
| NB038 | Y | N | Y | 248 | 115 | - |
| RA119 | Y | N | Y | 232 | 54 | - |
| VA105 | Y | N | N | - | - | - |
| 20F268 | Y | N | Y | 54 | 23 | - |

### Database-mining for marine DLOs

To further investigate any cryptic presence of DLOs in marine environments, publically available databases were mined. The BioMarKs, Global Ocean Sampling (GOS), Tara oceans marine metagenome, and Tara oceans V9 databases were all queried for the presence of DLO rRNA using the *Diphylleia*, KIVT02 and Å85 *Collodictyon* rRNA sequences. Further, the GOS and Tara oceans marine metagenome databases were queried with the 124 *Collodictyon* gene transcripts previously used to infer the phylogenetic placement of Diphyllatea (Zhao et al., 2012).

### Clustering, alignment construction and phylogenetic analyses

PacBio sequencing reads were split into their respective samples and barcodes removed using SMRT portal. Reads were subsequently filtered keeping a CCS read accuracy of 1.0. Reads lacking either the forward or reverse primer sequence were discarded. Reads were then clustered at 98% identity, sample dependent, utilizing the “-cluster-otus” command in Usearch v8.1 (Edgar, 2010; Edgar et al., 2011). The “-cluster-otus” command additionally removed possible chimeric reads from the dataset. Uncultured clones from the same locality, previously categorised as DLOs, acquired from NCBInr, were also clustered at the same identity. A 98% clustering identity was employed to allow for a long amplicon length, encompassing both conserved and variable regions. Additionally, we wanted to achieve a high diversity, possibly at a level lower than that of the species. Comparatively, a 97% clustering identity is normally applied to illumina amplicons when targeting variable rRNA regions (Wu et al., 2015). Clustered reads (OTUs) were then queried with blastn against a private database on CLC main workbench 7 (Qiagen) containing a broad selection of eukaryotic 18S rRNA taxa, including Diphyllatea (Årungen, *Diphylleia rotans* and KIVT02). OTU hits with an E-value of 0.0 to Diphyllatea were subsequently aligned to the previously constructed alignment using MAFFT “—add” with default parameters and manually refined with Mesquite v3.1 (Maddison and Maddison, 2017). Phylogenetic placement of the OTUs was checked using RAxML (method subsequent), with only those clustering with the Diphyllatea clade and constituting >1 read being kept for further analysis. Though, to check if a possible cryptic diversity was not disregarded, the method was repeated on reads with a CCS accuracy <1.0. After the removal of ambiguously aligned characters, using Gblocks with least stringent parameters (Talavera and Castresana, 2007), the final dataset consisted of 64 taxa and 3983 characters. The alignment (both masked and unmasked) has been made freely available through the authors ResearchGate pages (https://www.researchgate.net/home).

Maximum likelihood phylogenetic analyses were carried out using the GAMMA-GTR model in RAxML v8.0.26 (Stamatakis, 2006). The topology with the highest likelihood score of 100 heuristic searches was chosen. Bootstrap values were calculated from 500 pseudo-replicates. Bayesian inferences were performed using MrBayes v3.2.2 (Huelsenbeck and Ronquist, 2001), applying the GTR + GAMMA + Covarion model. Two independent runs, each with three cold and one heated Markov Chain Monte Carlo (MCMC) chains, were started from a random starting tree. The MCMC chains lasted for 40,000,000 generations with the tree sampled every 1000 generations. The posterior probabilities and mean marginal likelihood values of the trees were calculated after the burn-in phase, which was determined from the marginal likelihood scores of the initially sampled trees. The average split frequencies of the two runs were < 0.01, indicating the convergence of the MCMC chains.

To investigate any possible topological effect of inferring taxa with missing sequence data, a secondary alignment constituting the 18S rRNA was constructed (64 taxa and 1575 characters). This was inferred, as previous, and the topological congruence of the ingroup taxa was compared to that of the larger rRNA dataset using the Icong index (http://max2.ese.upsud.fr/icong/index.help.html) (de Vienne et al., 2007). Topologies were more congruent than expected by chance (Icong = 2.69 & P-value = 6.93e-12), rejecting any negative effect of inferring taxa with missing sequence data. As such, only the result for the larger rRNA analysis is presented (Fig. 3), with the 18S rRNA tree supplied as supplementary material (Supplementary Fig. 2)

**Fig. 1.**
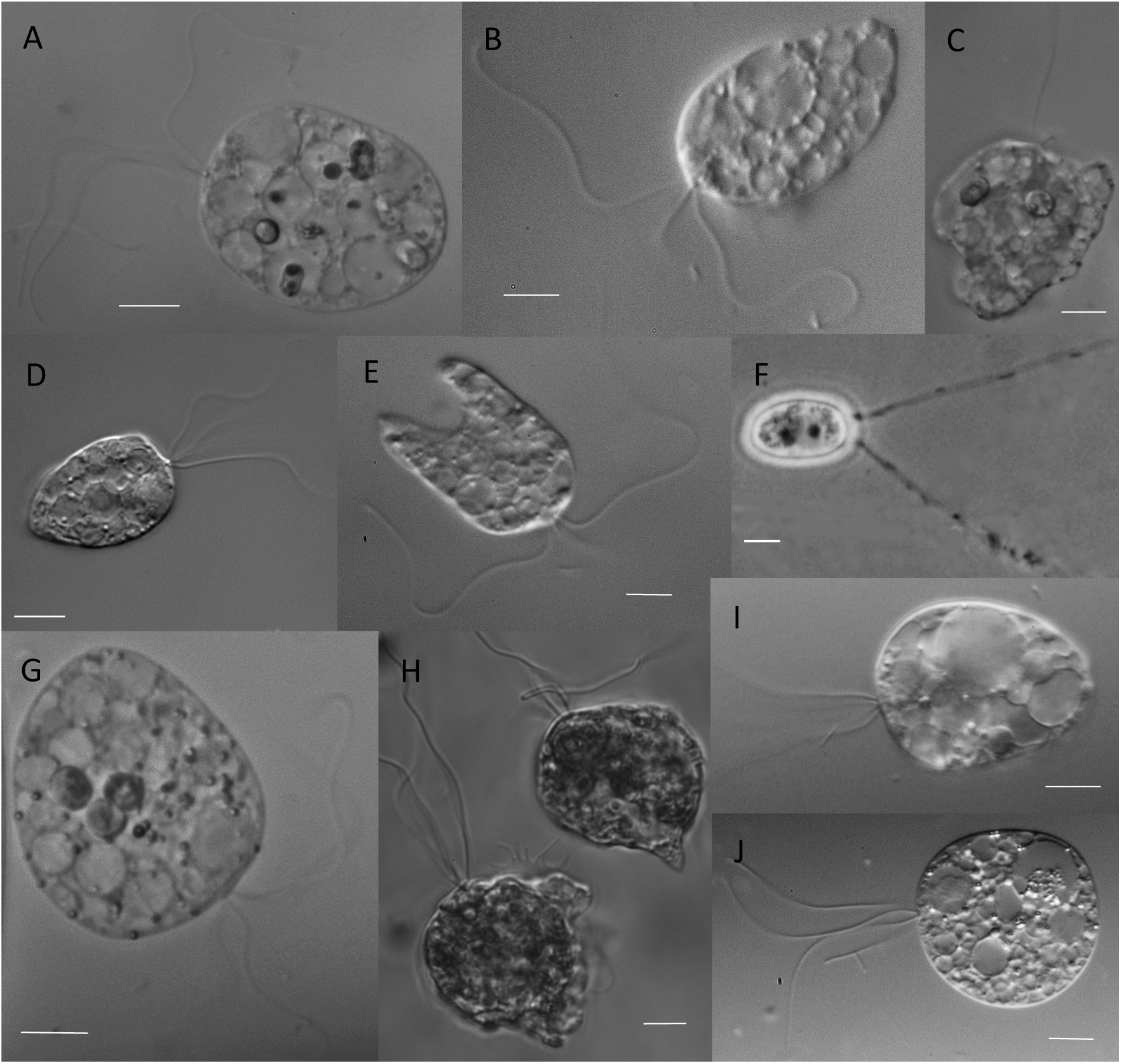
DIC Micrographs of newly cultured Diphyllatea-like organisms (Asian strains) and *C. triciliatum* (strain Å85). A) KIVTT01. B) KIINB. C) KIKNR01. D) KIVTT02. E) KIKNR02. F) Å85. G) KIVT04. H) Å85. I) Å85. J) KIVT01; scale bar 10 μm. The Diphy II is represented in H & I, whilst Diphy III is represented in A-E, G & J.

**Fig. 2.**
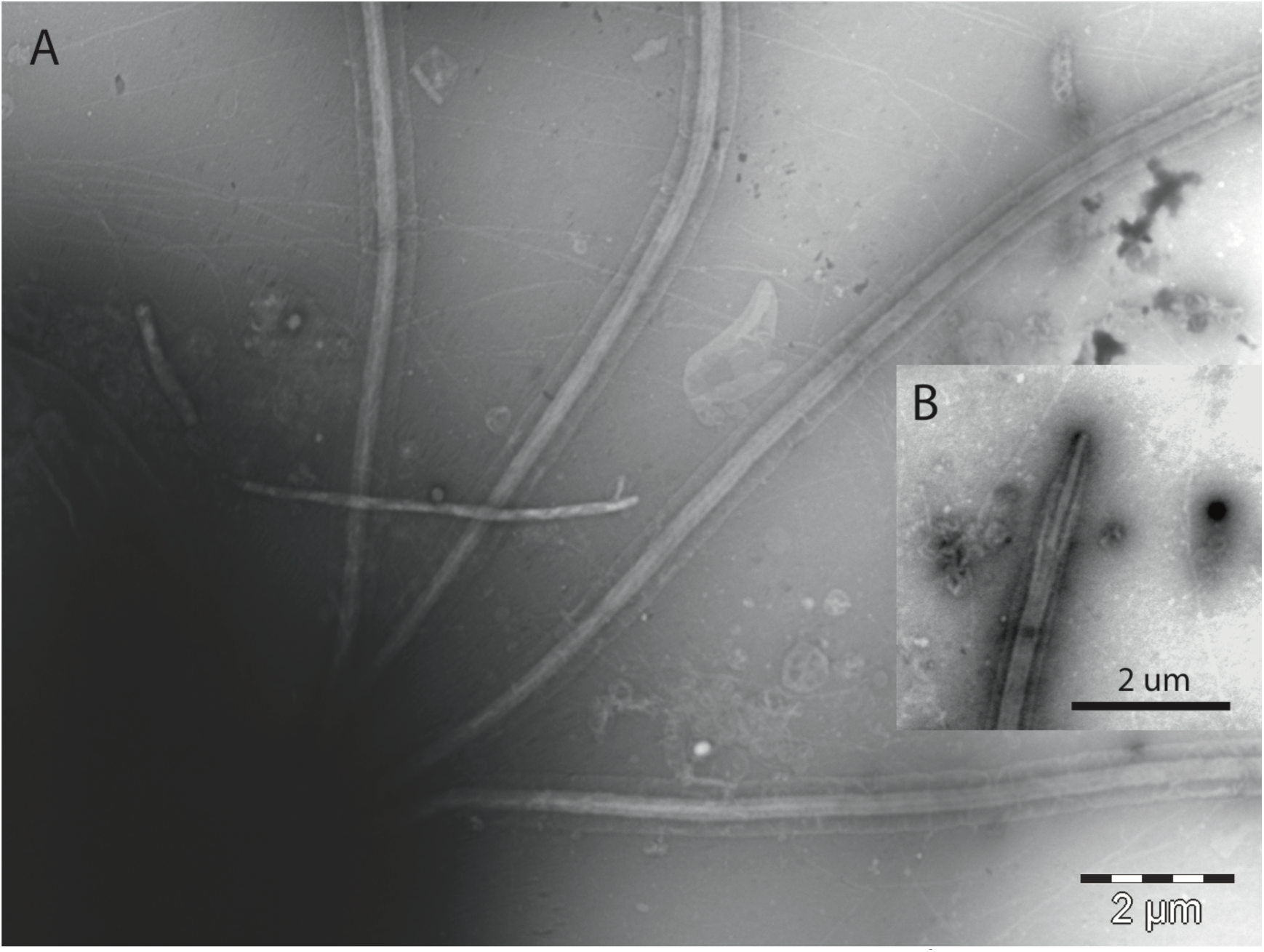
Electron micrographs of the flagella of *C. triciliatum* (strain Å85). (A) four flagella and the membranes of flagella; (B) tip of flagellum.

**Fig. 3.**
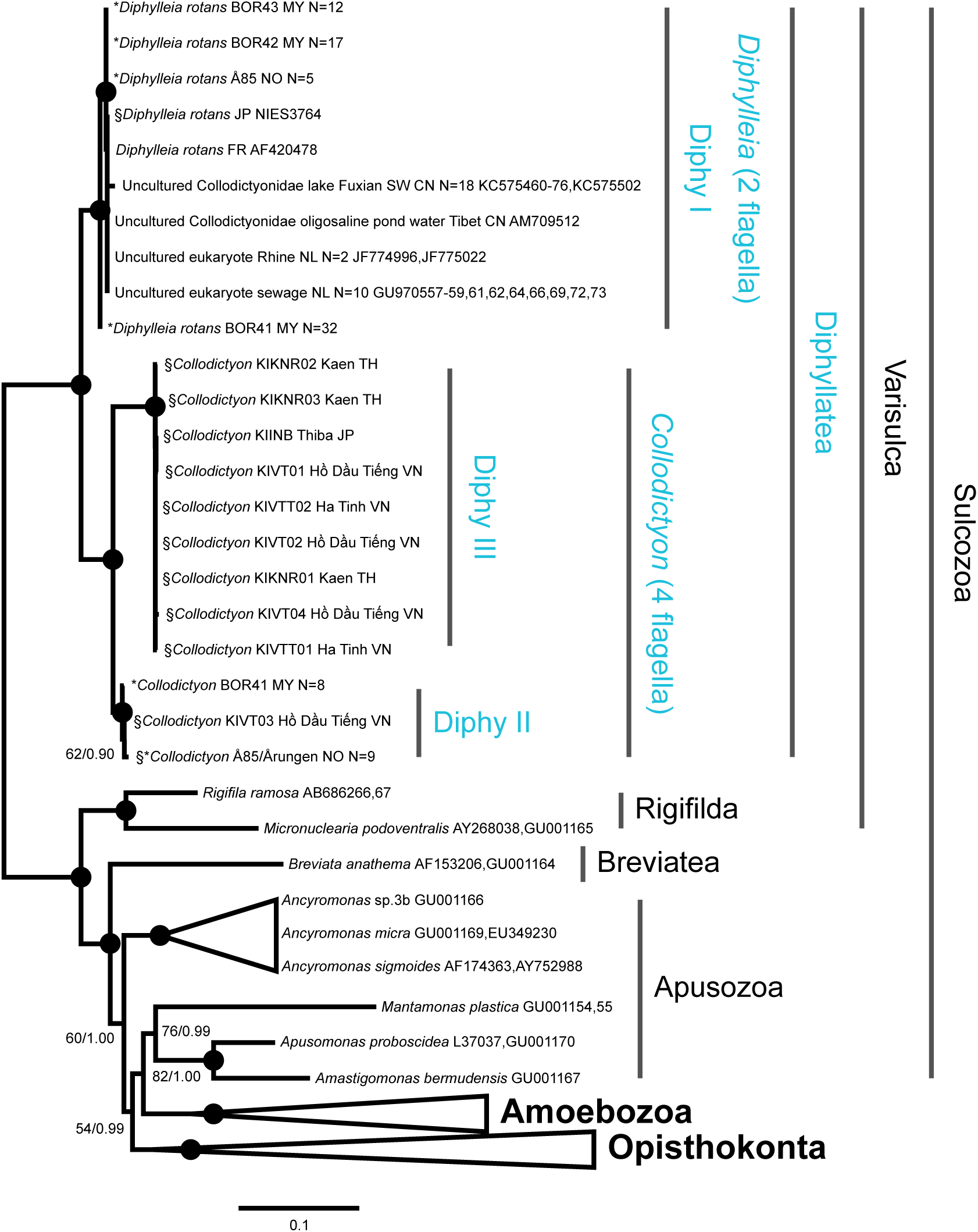
The rRNA phylogeny of Diphyllatea. The topology was reconstructed with the GAMMA-GTR model in RAxML v8.0.26. and inferred with 64 taxa and 3983 characters. The inference has been collapsed at varying taxonomic levels for easier visualisation, with blue representing the ingroup. The numbers on the internal nodes are ML bootstrap values (BP, inferred by RAxML v8.0.26. under then GAMMA-GTR model) and posterior probabilities (PP, inferred by MrBayes v3.2.2 under the GTR+GAMMA+Covarion model), ordered; RAxML / MrBayes. Black circles indicate BP > 90% and PP 1.00, values with BP < 50% are not shown. Asterisk (*) denotes environmental OTUs sequenced in this study, with “N” representing the number of reads included in each OTU. § depicts rRNA from cultured DLOs amplified in this study. Abbreviations for countries: CN = China, FR = France, JP = Japan, MY = Malaysia, NL = Netherlands, NO = Norway, TH = Thailand, and VN = Vietnam. See Supplementary Fig. 2 for 18S rRNA inference of Diphyllatea.

## Results and Discussion

### Cultured DLOs show a *Collodictyon* morphology

Diphyllatea is one of the earliest branching eukaryote lineages and as such of pivotal importance for reconstructing eukaryotic evolution. Still, virtually nothing is known about the higher order phylogenetic structure, diversity and geographic dispersal of this deep lineage. Here we address these issues by combining culturing, rRNA sequencing of new strains and targeted 18S rRNA PCR of natural samples sequenced with long-range high throughput PacBio RS technology.

Our survey of the diversity of Diphylleia generated 10 new strains from Japan, Thailand and Vietnam, which complemented the Norwegian *Collodictyon* strain already established in our lab (Table 1). Light microscope observations of all isolates showed a congruent morphology to *Collodictyon* by sharing an egg-or heart-like body and four isomorphic flagella (Fig. 1). All cultured strains were compared with the valid diagnoses of Diphyllatea and assigned to the morphospecies *C. triciliatum* (Carter, 1865). The swimming cells have a slow and relaxed movement, while rotating, driven by flagella (Fig. 1B, 1D, 1E, 1I, 1J and Supplementary Video 1). They have the ability to cling to the surface of the culture dish by a cytoplasmic veil and pseudopodia (i.e. the amoeboid property) within or from the sulcus (Fig. 1C, 1H and Supplementary Video 2). Observations of live cells show the central cytoplasm contains a few large or small vacuoles. Some of the vesicles engulf food particles (i.e. *Microcystis* strain CYA 43) at various stages of digestion (See Fig. 1A and 1G). All studied strains can form a ventral furrow or groove that extends dorsally dividing the cell into two parts (Fig. 1D and 1I), a non-permanent structure during the cell cycle. Emergence of a long groove may be exclusive to cells that are starving or initiating cell division. Furthermore, we uncovered thick-walled resting stages, cyst, with two long gelatinous filaments (Fig. 1F), similar to earlier descriptions of *Collodictyon* resting spores (Mischke, 1994), observed in the Å85 culture.

Electron microscopy of the negatively stained *Collodictyon* cells showed identical and smooth flagella lacking hairs or tomentum (Fig. 2). Further, the method confirmed that the periplast of *Collodictyon* is hyaline and even. Other sub-cellular ultrastructures were difficult to identify; the cell is highly fragile and easily disrupted in the electron microscopy fixation process (Klaveness, 1995).

Morphologically these isolates all seem to be strains of the *C. triciliatum* or closely related species, supporting earlier studies that Diphyllatea encompasses a limited species number (Carter, 1865; Francé, 1899; Massart, 1920; Brugerolle, 2006). Alternatively, a similar phenotype may represent multiple cryptic species with distinct genotypes (or ribotypes). We therefore investigated species diversity by sequencing rRNA, to reveal possible molecular differences between the strains.

### Diphyllatea is divided into three higher order groups (Diphy I – III)

The rRNA fragment of ∼6.3kb for all 11 *Collodictyon* cultures were successfully amplified, sequenced and assembled (Accession nrs MF039356-MF039367).

With the inclusion of *Diphylleia rotans* rRNA (MF039365) the Diphyllatea class was divided into three ribotype groupings based on sequence length and indels; the amplified fragment ranged from ∼5.8kb in *Diphylleia rotans*, here named Diphy I (calculated from genome sequence), ∼5.9kb in Å85 and KIVT03, here named Diphy II, to ∼6.3kb in KIINB, KIKNR01, KIKNR02, KIKNR03, KIVT01, KIVT02, KIVT04, KIVTT01 and KIVTT02, here named Diphy III (see alignment). As such, the three groups (Diphy I, II and III) had a 69-79% pairwise identity over the amplicon length, and a 73-86% pairwise identity for the 18S rRNA gene.

To investigate the diversity of the class further we designed new Diphyllatea-specific primer pairs and applied these to environmental DNA samples from varying habitats. Presence of 18S rRNA was confirmed in all environmental samples using the universal eukaryote primers NSF83-1528R. The Diphyllatea-specific primer pair Diphy257F-Diphy1881R, designed in this study, successfully amplified 18S rRNA from four environmental freshwater samples (Table 3). The primer pair with lower specificity to known Diphyllatea 18S rRNA, Diphy453F – 1528R, successfully amplified template from three of the remaining freshwater samples (Table 3).

Sequencing these environmental amplicons on a SMRTcell confirmed Diphy257F-Diphy1881R successfully amplified targeted 18s rRNA from Diphyllatea, whilst Diphy453F – 1528R was unsuccessful in amplifying DLO template. Pacbio CCS reads were filtered and clustered before being aligned with the Diphy I-III 18s-28s rRNA fragments previously amplified (see alignment and Fig 3). The freshwater samples BOR41 and Årungen both showed presence of Diphy I and II sequence. The BOR42 and BOR43 samples contained only Diphy I sequence data, though this is a likely result of limited sequencing depth. Interestingly, none of our new environmental 18S rRNA amplicons showed similarity to the Diphy III ribotype. No additional ribotypes were amplified that clustered external to the three Diphyllatea groupings already confirmed (Diphy I-III).

The new primers Diphy257F and Diphy1881R therefore provide a promising tool for future investigations of the Diphyllatea diversity. It should be noted, however, that even though Diphyllatea 18S rRNA was successfully amplified, the new primers also amplified a putative protein-coding gene (4-diphospocytidyl-2C-methyl-D-erythritol kinase) from a possible novel brackish Actinobacteria (MF039368) in two samples (BOR41 and BOR43). As these sequences were not 18S rRNA they were easily discarded.

### Diphyllatea unconfirmed in marine habitats

To date, Diphyllatea species have only been observed in freshwater environments, including an estuary of a freshwater river (Massart, 1920). As the new PCR primers successfully amplify Diphyllatea 18S rRNA from environmental samples, we used them to investigation a possible cryptic diversity in marine environments. However, while the marine DNA samples were of good quality and previously used for large surveys of protist diversity (Logares et al., 2014), we were unable to amplify Diphyllatea 18S rRNA (Table 3) using the Diphy257F-Diphy1881R primer pair. Applying less stringent PCR conditions (i.e. lowering annealing temperature from 55°C to 50°C) had no effect on this result. As previous, and in an attempt to amplify novel DLO template rRNA with lower sequence identity, the primer pair Diphy453F-1528R was used, successfully amplifying product from three of the four marine DNA samples (Table 3). However, once these environmental amplicons had been sequenced on a SMRTcell, filtered, and clustered, none of the 192 OTUs they represented had an affinity to Diphyllatea (Table 3).

Furthermore, in our search for marine DLO sequences, we complemented our PCR and sequencing approach by searching public sequence databases. Despite querying four of the largest sequence databases (BioMarKs, GOS, Tara oceans marine metagenome, and Tara oceans V9) for the presence of DLO rRNA and two databases (GOS and Tara oceans marine metagenome) with 124 *Collodictyon* gene transcripts (Zhao et al., 2012), we were unable to identify any marine Diphyllatea-like sequences. Instead, we identified 30 freshwater 18S rRNA sequences in the NCBInr database additional to the know *Diphylleia rotans* sequence from France (AF420478) and the uncultured Collodictyonidae sequence from Tibet (AM709512): 18 sequences (KC575460-76, KC575502) were from Lake Fuxian SW China, 2 sequences (JF774996, JF775022) from Rhine river water, Netherlands, and 10 sequences (GU970557-59, 61, 62, 64, 66, 69, 72, 73) from sewage water, Netherlands. All showed highest pairwise affinity to the Diphy I ribotype (Fig 3).

Altogether, our PCR amplification and database searches could only discover DLOs in freshwater, suggesting that Diphyllatea has a restricted environmental limit to freshwater habitats. Knowledge of Diphyllatea habitat preferences and distributions in environmental systems is still limited. Therefore, the development of targeted PCR approaches, presented here, can be useful in future studies on additional environmental samples.

### The phylogeny of Diphyllatea: the classes diversity

Using all generated and acquired sequences in phylogenetic reconstruction (Fig. 3) resulted in a monophyletic Diphyllatea grouping with full bootstrap support (BS) and posterior probability (PP). Further, Diphyllatea formed a fully supported (100 BS / 1.00 PP) clade with *Rigifila ramosa* and *Micronuclearia podoventralis* (order Rigifilda) to constitute the subphylum Variscula (Cavalier-Smith, 2013). As previous, Variscula was sister to *Breviata anthema* (class Breviatea; (Cavalier-Smith, 2013)), albeit with medium BS (60) and full PP support. The subphylum Apusozoa was recovered as a paraphyly, separated by the fully supported Amoebozoa and Opisthokonta (Cavalier-Smith, 2013). The result, albeit with limited character and taxon sampling, reduces support for grouping the sub-phyla Varisulca and Apusozoa within the phylum Sulcozoa (Cavalier-Smith, 2013; Zhao et al., 2013).

Sequences within Diphyllatea (Fig. 3) were divided into three clades, all fully supported. Two of these include the previously known genera *Diphylleia* and *Collodictyon*, here marked Diphy I and II respectively. Furthermore, and most importantly, the tree reveals a new Diphyllatea clade distinct from the previous two. The new clade, here named Diphy III, branches off as a sister clade to *Collodictyon* (Diphy II) with full support. All 11 cultured strains are placed within *Collodictyon* (Diphy II and III); two placing within Diphy II (Å85 and KIVT03), and the remaining nine within Diphy III (KIINB, KIKNR01, KIKNR02, KIKNR03, KIVT01, KIVT02, KIVT04, KIVTT01 and KIVTT02). The taxonomic rank Diphy I-III is not clear, but all groups contain higher diversity that earlier known, and several substructures that might constitute different sub-groups. To increase branching pattern resolution of the ingroup, the phylogeny was additionally inferred without outgroup taxa (Supp. Fig. 3). This inference was equivalent to that of Fig. 1, albeit with higher support for the observed branching patterns. The major difference being the separation of Diphy III strains into two supported clades. It should also be noted that Diphy III has a comparable basal branch to Diphy I and II, and therefore likely to be a clade at least on the same taxonomic level. The branch length is reflected in the sequence divergence between Diphy II and III: Only 79% sequence identity was shared over the rRNA length, increasing to 85% when considering the more conserved 18S rRNA region. Another distinct pattern in the tree, is the placement of *Diphylleia* BOR41 environmental OTU, which was placed as sister to Diphy I and excluded from this clade with almost full support (99 / 1.00), and 95 BS in Supp. Fig. 3, but still showed >98% pairwise identity to the other OTUs in Diphy I, suggesting *Diphylleia* likely constitutes several species (Fig. 3 & Supp. Fig. 3).

Mapping the morphology to the tree shows that the quadraflagellate forms branch together as two main monophyletic groups (i.e. Diphy II and III), implying that Diphyllatea as a group is deeply divided into two stems composed of quadraflagellate or biflagellate (Diphy I) forms. As the flagella of *Collodictyon* occupy the same position as the basal body in a pre-division stage of *Diphylleia* (Brugerolle et al., 2002), and the cyst stage has two long gelatinous filaments (Mischke, 1994), it could be hypothesised that the two forms represent different life-stages of the same species. However, no biflagellate stage was observed for our Diphy II and III cultures, with the phylogeny consistently separating the biflagellate *Diphylleia* from the two clades of quadraflagellate *Collodictyon*-like species. Hence, the morphological change in Diphyllatea has most likely occurred early in the history of the group. Because of the deep origin of Diphyllatea, this event represents one of the most ancient morphological innovations known among all eukaryotes. Existing sequence data suggests a single biflagellate clade, and as such may imply that the separation of *Diphylleia rotans* and *Sulcomonas lacustris,* based on cell size (Brugerolle, 2006) is unfounded. However, we are presently unable to exclude a fourth Diphyllatea species despite lacking amplicon and database evidence.

### A global distribution of Diphyllatea

The substantial increase in Diphyllatea sequence data presented, allows, for the first time, conclusions to be drawn as to the extent and ecological role of the class and genera therein.

The Diphy I clade (Fig. 3), representing the biflagellate cell-type, constituted sequences from Borneo, China, France, Japan, Netherlands and Norway, including the previously reported ribotypes from Tibet (China) and Clermont-Ferrand (France) (Brugerolle et al., 2002; Wu et al., 2009). In addition to the localities above, Diphylleia has also be reported in Saudi Arabia (Mohamed and Al-Shehri, 2013), suggesting a possible global distribution of the genus.

The Diphy II clade constituted sequences from Borneo and Norway, from both culture and amplicon. The Diphy III clade, in contrast, constituted only cultured sequence data from Asia (Japan, Thailand and Vietnam) with no environmental DLO amplicons having an affinity to this clade. *Collodictyon*, the quadraflagellate form (represented in the Diphy II and III clades; Fig. 3), has been previously described from the island of Bombay and later in central Europe, Spain and Norway (Carter, 1865; Rhodes, 1917; Klaveness, 1995; Sánchez et al., 1998). The quadraflagellate morphotype has additionally been reported in North America (Carter, 1865; Rhodes, 1917; Lackey, 1942), and more recently South America; from multiple freshwater localities in Uruguay: La Oriental, Maldonado (34°34’S 55°15’W), Tala, Canelones (34°20’S 55°45’W), and Picada Varela, San José River, San José (34°19’S 56°42’W). Accompanying video is available through https://www.youtube.com/ (uuvb3eUZUQ8, AsY8s-HnTMQ, M8tAf3KoDQM and k88LsRcEXmg). As only morphological data is presented, in the reports above, we are unable to establish if these morphotypes represent Diphy II and/or III, however it does confirm a global distribution of the *Collodictyon* morphotype and accordingly the Diphyllatea class.

Interestingly, all environmental sequence data for DLOs deposited in public databases were only related to the Diphy I clade. The reason for this pattern is unclear but unlikely a result of the PCR primers used, which showed no mismatch against a Diphy II and III 18S rRNA target. It may rather indicate higher abundance of Diphy I in the sampled localities or could reflect different habitat preferences among the three Diphy groups.

Despite the cryptic diversity of Diphyllatea in natural communities, its ecological importance should not be ignored. There are a variety of food sources for DLO strains (Mischke, 1994; Klaveness, 1995; Brugerolle et al., 2002; Brugerolle, 2006), and they may have a broader selection of prey in their natural environments (Francé, 1899; Bělař, 1926; Wawrik, 1973). DLOs can consume nano-and picoplankton (e.g. cryptomonads, euglenoids, green algae and cyanobacteria) and are therefore considered to hold a crucial position in the food web. Here, the study of freshwater Diphyllatea offered a first glimpse of its diversity and distribution. As more DLO sequences are identified in future cultural and environmental surveys, our insight into the living habitat and ecological role of Diphyllatea will be further elucidated.

### PacBio SMRT sequencing of targeted Diphyllatea amplicons

The PacBio RS sequencing platform has been previously used to study 16s (Fichot and Norman, 2013; Mosher et al., 2013) and more recently rRNA amplicons (Jones and Kustka, 2017; Tedersoo et al., 2017). Though, in contrast to these studies, that wanted to answer a broader diversity question, our goal was a targeted 18S rRNA approach with PacBio RS replacing traditional cloning methods. Our result demonstrates PacBio RS as an efficient and economical alternative to the traditional cloning and sanger method for sequencing long rRNA amplicons (Edgcomb et al., 2011; Thomas et al., 2012; Bachy et al., 2013). A single SMRT cell gave 6,310 total reads of insert (Supplementary Table. 2), which constituted 1,741 high quality sequences (CCS = 1) and 281 OTUs. To achieve a comparable number of sequences via cloning and Sanger sequencing would be a tedious exercise, at an approximate cost 30x higher than that of our PacBio RS method (based on CCS = 1 result).

The major advantage of PacBio RS for the study eukaryotic of diversity is read length, which allows for higher phylogenetic resolution. Additionally, long reads allow short read amplicon datasets from contrasting rRNA regions to be “scaffolded” and inferred in parallel, further increasing resolution. However, PacBio RS does have a high error rate, ∼15% with the P4-C2 chemistry, that is overcome by CCS, and further accounted for with a sequence analysis pipeline. Our results demonstrate an analysis pipeline as essential for the removal of both sequencing errors and chimeras, that increase proportional with amplicon length (Laver et al., 2016; Tedersoo et al., 2017), which can give an overestimation of diversity. Of the 1,741 high quality sequences, 186 (11.44%) were identified as chimeric using Uchime (see Supplementary Table 2). The high level of chimeric sequences identified was surprising, with only a 1-2% chimera level previously reported (Koren et al., 2012; Fichot and Norman, 2013) by “misligation” of SMRTbell adaptors in the PacBio library preparation. The observed chimera level is therefore attributed to PCR artefacts; It has been proposed that >45% of reads in some datasets are chimeric (Huber et al., 2004; Ashelford et al., 2006; Laver et al., 2016), with experiments showing that >30% of chimeras can be attributed to PCR (Wang and Wang, 1997). Formation of chimeras increases with amplicon length, where template switching is prone to chimeric product formation (Laver et al., 2016), as such filters to identify and remove long read amplicon chimeras (i.e. Uchime) are paramount. It is difficult to ascertain if the observed chimeras are a result of the original 18S rRNA amplification or the subsequent PCR to attach symmetric PacBio barcodes. It is possible, however to reduce the former by decreasing amplification cycles (Smyth et al., 2010), and eradicate the latter by barcode ligation or the sequencing of separate samples, instead of multiplexing. It has been previously reported that Uchime fails to identify all long read amplicon chimeras (Fichot and Norman, 2013), though we found no evidence supporting this in our dataset.

However, and as with all sequencing platforms, PacBio will improve with technological and chemical developments, allowing for longer and more accurate reads with higher output (Fichot and Norman, 2013), with the recent release of PacBio Sequel confirming (Hebert et al., 2017; Tedersoo et al., 2017). Further, understanding PacBio biases will allow for improved bioinformatic pipelines and as such phylogenetic inferences (Fichot and Norman, 2013). It should be noted that Oxford Nanopore sequencing platform has recently been applied to 16S amplicons (Ma et al., 2017), and as such can offer an alternative for the study of eukaryotic diversity with long reads.

### Conclusions and future perspectives

Our results suggest that combining culture methods with environmental PCR is invaluable for understanding the diversity and distribution of protist lineages, in particular Diphyllatea, and their ecological importance in aquatic systems. Here we provide the tools to uncover the true diversity of this class. In this study, the application of DLO culturing techniques and the sequencing of the partial rRNA operon from multiple strains, reveals two clades, Diphy II and III with quadraflagellate morphology. The inference of DLO cultured sequences with that of database orthologues and environmental amplicons infers a greater Diphyllatea diversity than previously known, recovering three clearly phylogenetically separated clades (Diphy I, II, and III). Biflagellate and quadraflagellate Diphyllatea species were separated, suggesting morphological innovation has occurred deep within the evolution of the class. Further, we show the Diphyllatea class to have a global distribution limited to freshwater. To further understand the evolution of Diphyllatea, the ancestral form, a possible genome duplication, and its relationship within Sulcozoa, the genomes of multiple species are essential. For this reasoning, we are presently completing the annotation of genomes from each of the three Diphyllatea clades, in addition to that of *Rigifila ramosa* (Rigifilda) and a new Breviatea species.

Lastly, our study shows the capacity of PacBio RS when employing a targeted approach for increasing phylogenetic resolution of an enigmatic and deeply diverging protists. Although caution needs to be observed when analysing reads, to avoid an overestimation of diversity, the platform offers major economical and efficiency gains over traditional cloning and Sanger sequencing methods, something that will improve with technological advances.

## Acknowledgements

We would like to thank Abel (UiO) for providing computing resources, in particular projects nn9244k and nn9404k. We thank the Norwegian Sequencing Centre (NSC) for providing advice related to PacBio sequencing and analysis. We are grateful to Dr. David Bass and Dr. Bente Edvardsen, representing Biomarks, for providing environmental DNA samples. The Norwegian Institute for Water research (NIVA) are acknowledged for providing the *Microcystis* strain CYA 43. We thank Katherine Schou and Dr. Anders Krabberød (UiO) for laboratory support and discussion in formulating this paper. Thanks, are given to Dr. Takashi Shiratori and Dr. Ken Ishida from the University of Tsukuba for providing the *D. rotans* strain for a separate genomic study. This work has been supported by research grants from the Norwegian Research Council to R.J.S.O (project 230868) and the University of Oslo to K.S.-T. and D.K.

## References

Al-Bulushi, I.M., Bani-Uraba, M.S., Guizani, N.S., Al-Khusaibi, M.K., and Al-Sadi, A.M. (2017) Illumina MiSeq sequencing analysis of fungal diversity in stored dates. BMC Microbiology 17: 72.

Asemaninejad, A., Weerasuriya, N., Gloor, G.B., Lindo, Z., and Thorn, R.G. (2016) New Primers for Discovering Fungal Diversity Using Nuclear Large Ribosomal DNA. PLOS ONE 11: e0159043.

Ashelford, K.E., Chuzhanova, N.A., Fry, J.C., Jones, A.J., and Weightman, A.J. (2006) New Screening Software Shows that Most Recent Large 16S rRNA Gene Clone Libraries Contain Chimeras. Applied and Environmental Microbiology 72: 5734–5741.

Bachy, C., Dolan, J.R., López-García, P., Deschamps, P., and Moreira, D. (2013) Accuracy of protist diversity assessments: morphology compared with cloning and direct pyrosequencing of 18S rRNA genes and ITS regions using the conspicuous tintinnid ciliates as a case study. The ISME Journal 7: 244–255.

Bass, D., and Cavalier-Smith, T. (2004) Phylum-specific environmental DNA analysis reveals remarkably high global biodiversity of Cercozoa (Protozoa). International Journal of Systematic and Evolutionary Microbiology 54: 2393–2404.

Bělař, K.I. (1926) Der Formwechsel der Protistenkerne: Eine vergleichend-morphologische Studie: Ergebn. u. Fortschr. der Zoologie.

Bo-Ra, K., Shin-ichi, N., Baik-Ho, K., and Myung-Soo, H. (2006) Grazing and growth of the heterotrophic flagellate Diphylleia rotans on the cyanobacterium Microcystis aeruginosa. Aquatic Microbial Ecology 45: 163–170.

Bradley, I.M., Pinto, A.J., and Guest, J.S. (2016) Design and Evaluation of Illumina MiSeq-Compatible, 18S rRNA Gene-Specific Primers for Improved Characterization of Mixed Phototrophic Communities. Applied and Environmental Microbiology 82: 5878–5891.

Bråte, J., Klaveness, D., Rygh, T., Jakobsen, K.S., and Shalchian-Tabrizi, K. (2010) Telonemia-specific environmental 18S rDNA PCR reveals unknown diversity and multiple marine-freshwater colonizations. BMC Microbiology 10: 168.

Brown, M.W., Sharpe, S.C., Silberman, J.D., Heiss, A.A., Lang, B.F., Simpson, A.G.B., and Roger, A.J. (2013) Phylogenomics demonstrates that breviate flagellates are related to opisthokonts and apusomonads. Proceedings of the Royal Society B: Biological Sciences 280: 20131755.

Brugerolle, G. (2006) Description of a New Freshwater Heterotrophic Flagellate Sulcomonas lacustris Affiliated to the Collodictyonids. Acta Protozoologica 45: 175–182.

Brugerolle, G., and Patterson, D.J. (1990) A cytological study of Aulacomonas submarina Skuja 1939, a heterotrophic flagellate with a novel ultrastructural identity. European Journal of Protistology 25: 191–199.

Brugerolle, G., Bricheux, G., Philippe, H., and Coffea, G. (2002) Collodictyon triciliatum and Diphylleia rotans (= Aulacomonas submarina) form a new family of flagellates (Collodictyonidae) with tubular mitochondrial cristae that is phylogenetically distant from other flagellate groups. Protist 153: 59–70.

Burki, F., Kaplan, M., Tikhonenkov, D.V., Zlatogursky, V., Minh, B.Q., Radaykina, L.V. et al. (2016) Untangling the early diversification of eukaryotes: a phylogenomic study of the evolutionary origins of Centrohelida, Haptophyta and Cryptista. Proceedings of the Royal Society B: Biological Sciences 283: 20152802.

Carter, H.J. (1865) XXXII. — On the fresh- and salt-water Rhizopoda of England and India. In Annals and Magazine of Natural History: Taylor & Francis, pp. 277–293.

Cavalier-Smith, T. (2013) Early evolution of eukaryote feeding modes, cell structural diversity, and classification of the protozoan phyla Loukozoa, Sulcozoa, and Choanozoa. European Journal of Protistology 49: 115–178.

Cavalier-Smith, T., and Chao, E.E. (2010) Phylogeny and Evolution of Apusomonadida (Protozoa: Apusozoa): New Genera and Species. Protist 161: 549–576.

de Vargas, C., Audic, S., Henry, N., Decelle, J., Mahé, F., Logares, R. et al. (2015) Eukaryotic plankton diversity in the sunlit ocean. Science 348.

de Vienne, D.M., Giraud, T., and Martin, O.C. (2007) A congruence index for testing topological similarity between trees. Bioinformatics 23: 3119–3124.

Doyle, J.J., and Doyle, J.L. (1987) A rapid DNA isolation procedure for small quantities of fresh leaf tissue. Phytochemistry Bulletin, Botanical Society of America 19: 11–15.

Edgar, R.C. (2010) Search and clustering orders of magnitude faster than BLAST. Bioinformatics 26: 2460–2461.

Edgar, R.C., Haas, B.J., Clemente, J.C., Quince, C., and Knight, R. (2011) UCHIME improves sensitivity and speed of chimera detection. Bioinformatics 27.

Edgcomb, V., Orsi, W., Bunge, J., Jeon, S., Christen, R., Leslin, C. et al. (2011) Protistan microbial observatory in the Cariaco Basin, Caribbean. I. Pyrosequencing vs Sanger insights into species richness. ISME J 5: 1344–1356.

Fichot, E.B., and Norman, R.S. (2013) Microbial phylogenetic profiling with the Pacific Biosciences sequencing platform. Microbiome 1: 10.

Francé, R. (1899) A Collodictyon triciliatum Cart. Szervezete (Über den Organismus Collodictyon triciliatum Cart.). In Termeszetrai füzetek. Budapest: Tabla, pp. 1–26.

Gadberry, M.D., Malcomber, S.T., Doust, A.N., and Kellogg, E.A. (2005) Primaclade-a flexible tool to find conserved PCR primers across multiple species. Bioinformatics 21: 1263–1264.

Gordon, D., Abajian, C., and Green, P. (1998) Consed: A graphical tool for sequence finishing. Genome Research 8: 195–202.

Guillard, R.R.L., and Lorenzen, C.J. (1972) YELLOW-GREEN ALGAE WITH CHLOROPHYLLIDE C1,2. Journal of Phycology 8: 10–14.

Hebert, P.D., Braukmann, T.W., Prosser, S.W., Ratnasingham, S., deWaard, J.R., Ivanova, N.V. et al. (2017) A Sequel to Sanger: Amplicon Sequencing That Scales. bioRxiv.

Hendriks, L., Goris, A., Neefs, J.M., Van de Peer, Y., Hennebert, G., and Dewachter, R. (1989) The Nucleotide-Sequence of the Small Ribosomal-Subunit RNA of the Yeast Candida albicans and the Evolutionary Position of the Fungi among the Eukaryotes. Systematic and Applied Microbiology 12: 223–229.

Hoang, N.V., Furtado, A., Mason, P.J., Marquardt, A., Kasirajan, L., Thirugnanasambandam, P.P. et al. (2017) A survey of the complex transcriptome from the highly polyploid sugarcane genome using full-length isoform sequencing and de novo assembly from short read sequencing. BMC Genomics 18: 395.

Huber, T., Faulkner, G., and Hugenholtz, P. (2004) Bellerophon: a program to detect chimeric sequences in multiple sequence alignments. Bioinformatics 20: 2317–2319.

Huelsenbeck, J., and Ronquist, F. (2001) MrBayes: Bayesian inference of phylogenetic trees. Bioinformatics 17: 754–755.

Hugerth, L.W., Muller, E.E.L., Hu, Y.O.O., Lebrun, L.A.M., Roume, H., Lundin, D. et al. (2014) Systematic Design of 18S rRNA Gene Primers for Determining Eukaryotic Diversity in Microbial Consortia. PLOS ONE 9: e95567.

Jones, B.M., and Kustka, A.B. (2017) A quantitative SMRT cell sequencing method for ribosomal amplicons. Journal of Microbiological Methods 135: 77–84.

Katoh, K., and Standley, D.M. (2013) MAFFT Multiple Sequence Alignment Software Version 7: Improvements in Performance and Usability. Molecular Biology and Evolution 30: 772–780.

Kibbe, W.A. (2007) OligoCalc: an online oligonucleotide properties calculator. Nucleic Acids Research 35: W43–W46.

Kimura, B., and Ishida, Y. (1985) Photophagotrophy in Uroglena americana, Chrysophyceae. Japanese Journal of Limnology (Rikusuigaku Zasshi) 46: 315–318.

Klaveness, D. (1995) Collodictyon triciliatum H.J. Carter (1865)-a Common but Fixation-sensitive Algivorous Flagellate from the Limnopelagial. Nordic Journal of Freshwater Research 70: 3–11.

Koren, S., Schatz, M.C., Walenz, B.P., Martin, J., Howard, J.T., Ganapathy, G. et al. (2012) Hybrid error correction and de novo assembly of single-molecule sequencing reads. Nat Biotech 30: 693–700.

Kuo, R.I., Tseng, E., Eory, L., Paton, I.R., Archibald, A.L., and Burt, D.W. (2017) Normalized long read RNA sequencing in chicken reveals transcriptome complexity similar to human. BMC Genomics 18: 323.

Lackey, J.B. (1942) The Plankton Algae and Protozoa of Two Tennessee Rivers. The American Midland Naturalist 27: 191–202.

Laver, T.W., Caswell, R.C., Moore, K.A., Poschmann, J., Johnson, M.B., Owens, M.M. et al. (2016) Pitfalls of haplotype phasing from amplicon-based long-read sequencing. Scientific Reports 6: 21746.

Logares, R., Audic, S., Bass, D., Bittner, L., Boutte, C., Christen, R. et al. (2014) Patterns of Rare and Abundant Marine Microbial Eukaryotes. Current Biology 24: 813–821.

Ma, X., Stachler, E., and Bibby, K. (2017) Evaluation of Oxford Nanopore MinION Sequencing for 16S rRNA Microbiome Characterization. bioRxiv.

Maddison, W.P., and Maddison, D.R. (2017) Mesquite: a modular system for evolutionary analysis. Version 3.1 http://mesquiteproject.org/. In.

Massart, J. (1920) Recherches sur les organismes inférieurs. VIII.-sur la motilité des flagellates. Académie Royale d’e Belgique Bulletin de la Classe des Sciences 5: 116–141.

Medlin, L., Elwood, H., Stickel, S., and Sogen, M. (1988) The characterization of enzymatically amplified eukaryotic 16S-like rRNA-coding regions. Gene 71: 491–499.

Mischke, U. (1994) Influence of food quality and quantity on ingestion and growth rates of three omnivorous heterotrophic flagellates. Marine Microbial Food Webs 8: 125–143.

Mohamed, Z.A., and Al-Shehri, A.M. (2013) Grazing on Microcystis aeruginosa and degradation of microcystins by the heterotrophic flagellate Diphylleia rotans. Ecotoxicology and Environmental Safety 96: 48–52.

Mosher, J.J., Bernberg, E.L., Shevchenko, O., Kan, J., and Kaplan, L.A. (2013) Efficacy of a 3rd generation high-throughput sequencing platform for analyses of 16S rRNA genes from environmental samples. Journal of Microbiological Methods 95: 175–181.

Mueller, R.C., Gallegos-Graves, L.V., and Kuske, C.R. (2016) A new fungal large subunit ribosomal RNA primer for high-throughput sequencing surveys. FEMS Microbiology Ecology 92: fiv153–fiv153.

Orr, R.J.S., Rombauts, S., Van de Peer, Y., and Shalchian-Tabrizi, K. (2017a) Draft Genome Sequences of Two Unclassified Bacteria, Sphingomonas sp. Strains IBVSS1 and IBVSS2, Isolated from Environmental Samples. Genome Announcements 5.

Orr, R.J.S., Rombauts, S., Van de Peer, Y., and Shalchian-Tabrizi, K. (2017b) Draft Genome Sequences of Two Unclassified Chitinophagaceae Bacteria, IBVUCB1 and IBVUCB2, Isolated from Environmental Samples. Genome Announcements 5.

Potvin, M., and Lovejoy, C. (2009) PCR-Based Diversity Estimates of Artificial and Environmental 18S rRNA Gene Libraries. Journal of Eukaryotic Microbiology 56: 174–181.

Rhodes, R.C. (1917) Binary Fission in Collodictyon tricilliatum. Berkeley, California: University of California.

Ruggiero, M.A., Gordon, D.P., Orrell, T.M., Bailly, N., Bourgoin, T., Brusca, R.C. et al. (2015) A Higher Level Classification of All Living Organisms. PLOS ONE 10: e0119248.

Sánchez, J.C., Cobelas, M. Á., and Sanjurjo, M.A. (1998) Lista florística y bibliográfica de los clorófitos (Chlorophyta) de la Península Ibérica, Islas Baleares e Islas Canarias. the University of California: Asociación Española de Limnología.

Schmitt, I., Crespo, A., Divakar, P.K., Fankhauser, J.D., Herman-Sackett, E., Kalb, K. et al. (2009) New primers for promising single-copy genes in fungal phylogenetics and systematics. Persoonia 23: 35–40.

Smyth, R.P., Schlub, T.E., Grimm, A., Venturi, V., Chopra, A., Mallal, S. et al. (2010) Reducing chimera formation during PCR amplification to ensure accurate genotyping. Gene 469: 45–51.

Stamatakis, A. (2006) RAxML-VI-HPC: maximum likelihood-based phylogenetic analyses with thousands of taxa and mixed models. Bioinformatics 22: 2688–2690.

Stanier, R.Y., Kunisawa, R., Mandel, M., and Cohen-Bazire, G. (1971) Purification and properties of unicellular blue-green algae (order Chroococcales). Bacteriological Reviews 35: 171–205.

Talavera, G., and Castresana, J. (2007) Improvement of phylogenies after removing divergent and ambiguously aligned blocks from protein sequence alignments. Syst Biol 56: 564–577.

Taylor, D.L., Walters, W.A., Lennon, N.J., Bochicchio, J., Krohn, A., Caporaso, J.G., and Pennanen, T. (2016) Accurate Estimation of Fungal Diversity and Abundance through Improved Lineage-Specific Primers Optimized for Illumina Amplicon Sequencing. Applied and Environmental Microbiology 82: 7217–7226.

Tedersoo, L., Tooming-Klunderud, A., and Anslan, S. (2017) PacBio metabarcoding of Fungi and other eukaryotes: errors, biases and perspectives. New Phytologist.

Thomas, M.C., Selinger, L.B., and Inglis, G.D. (2012) Seasonal Diversity of Planktonic Protists in Southwestern Alberta Rivers over a 1-Year Period as Revealed by Terminal Restriction Fragment Length Polymorphism and 18S rRNA Gene Library Analyses. Applied and Environmental Microbiology 78: 5653–5660.

Wang, G.C., and Wang, Y. (1997) Frequency of formation of chimeric molecules as a consequence of PCR coamplification of 16S rRNA genes from mixed bacterial genomes. Applied and Environmental Microbiology 63: 4645–4650.

Wawrik, F. (1973) Collodictyon Studien. Arch Protistenkunde 115: 353–356.

Wu, L., Wen, C., Qin, Y., Yin, H., Tu, Q., Van Nostrand, J.D. et al. (2015) Phasing amplicon sequencing on Illumina Miseq for robust environmental microbial community analysis. BMC Microbiology 15: 125.

Wu, Q.L., Chatzinotas, A., Wang, J., and Boenigk, J. (2009) Genetic Diversity of Eukaryotic Plankton Assemblages in Eastern Tibetan Lakes Differing by their Salinity and Altitude. Microbial ecology 58: 569–581.

Zhao, S., Shalchian-Tabrizi, K., and Klaveness, D. (2013) Sulcozoa revealed as a paraphyletic group in mitochondrial phylogenomics. Molecular Phylogenetics and Evolution 69: 462–468.

Zhao, S., Burki, F., Bråte, J., Keeling, P.J., Klaveness, D., and Shalchian-Tabrizi, K. (2012) Collodictyon—An Ancient Lineage in the Tree of Eukaryotes. Molecular Biology and Evolution 29: 1557–1568.

